# A life history model of indeterminate growth, somatic maintenance, and negative senescence

**DOI:** 10.64898/2026.08.24.746687

**Authors:** Arttu Soukainen, Piret Avila

**Affiliations:** Faculty of Biological and Environmental Sciences, University of Helsinki, 00014 Helsinki, Finland

**Keywords:** life history evolution, negative senescence, optimal control theory, ageing, disposable soma theory, indeterminate growth

## Abstract

Some organisms exhibit declining mortality and increasing fecundity following sexual maturity, a demographic pattern known as negative senescence. According to life history theory, ageing occurs because resources are preferentially allocated to reproduction over somatic maintenance. Models connecting indeterminate growth to negative senescence exist, but none integrate somatic maintenance as a competing allocation decision alongside growth and reproduction. We formulate a life history model in which an individual allocates energy among reproduction, somatic growth, and somatic maintenance and mortality rate depends on both body size and somatic damage. We show that negative actuarial senescence, whereby mortality declines with age, occurs when the proportional change in reproductive value exceeds the proportional change in fitness returns from current investments into reproduction and soma. We derive the necessary conditions for an uninvadable allocation strategy using invasion analysis and Pontryagin’s maximum principle, and examine biologically relevant cases numerically. We show that both negative senescence and indeterminate growth arise together as uninvadable outcomes even when maintenance competes for the same resources as growth and reproduction. We show that both diminishing returns to reproduction and diminishing returns to growth can give rise to negative senescence. These results extend the disposable soma theory to organisms with indeterminate growth, in which mortality decreases with size, and identify key mechanisms for the empirically observed association between indeterminate growth and non-senescent demographic trajectories.

## 1 Introduction

Senescence, commonly understood as the age-related decline in survival and reproduction, is widespread in the tree of life. Yet ageing patterns show remarkable variation and sustained increases in fertility and decreases in mortality after maturity have been documented across a diverse range of taxa, particularly among species with indeterminate growth (continued growth after sexual maturity) (Finch, 1994; Jones et al., 2014; Shefferson et al., 2017). Despite this empirical evidence, classical evolutionary theories of ageing have focused predominantly on explaining why senescence occurs (Medawar, 1952; Williams, 1957; Hamilton, 1966; Kirkwood, 1977), and the conditions under which vital rates can improve with age have received comparatively little theoretical attention (but see Vaupel et al., 2004; Baudisch, 2008).

Senescence is the age-related decline in demographic performance, expressed through two vital rates: an increase in mortality *µ*(*t*) (actuarial senescence) and a decline in fecundity *b*(*t*) (reproductive senescence), after reproductive maturity (first reproduction onwards). However, the two rates need not change in the same direction, or at the same proportional rate, as individuals age. For this reason, it has been proposed that a useful definition is that senescence occurs at age *t* when the proportional rate of increase in mortality, 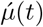, exceeds that of fecundity, 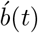, i.e. 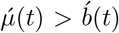 (Baudisch, 2008, see caption of Fig. 1 for further details about this definition). We endorse this classification here, and here we call the complementary case non-senescence, whereby 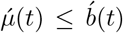. Within non-senescence, we distinguish two cases. *Negative senescence* whereby mortality is strictly declining and fecundity is stable or increasing. All remaining cases of non-senescence, for instance, a slight increase in mortality offset by a substantial increase in fecundity, we term *negligible senescence*. Figure 1 summarises this classification.

**Figure 1.**
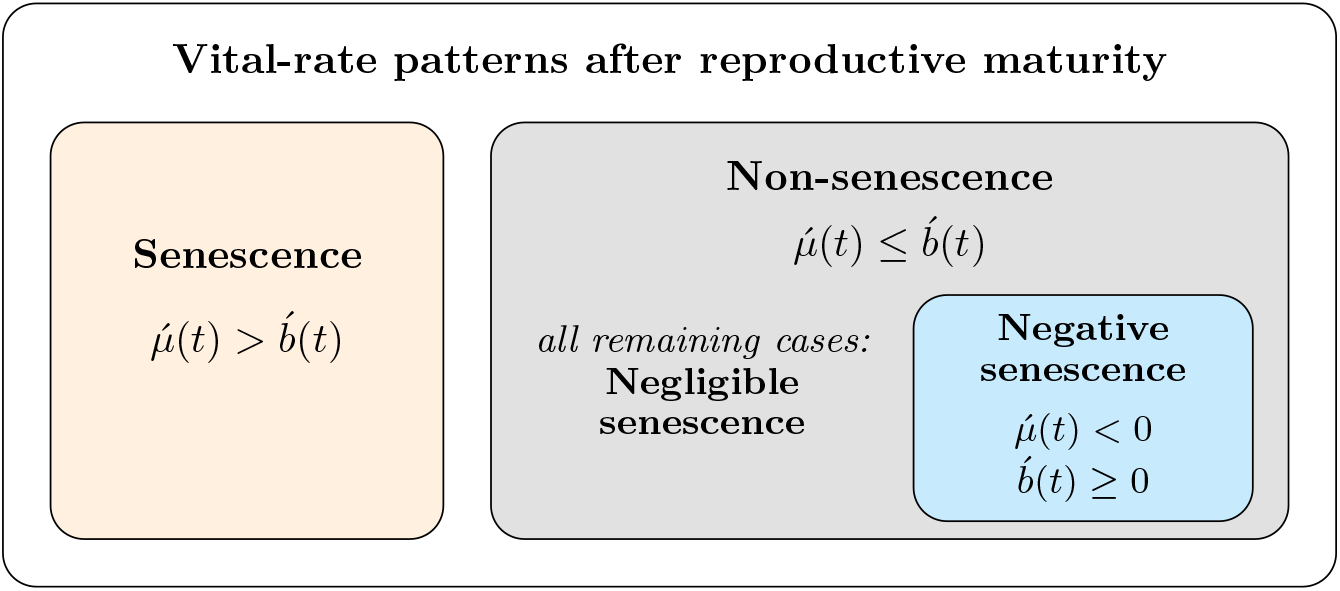
Classification of age-specific vital rate patterns after reproductive maturity in terms of fecundity *b*(*t*) and mortality *µ*(*t*). Here, 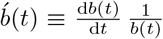 and 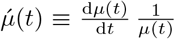 denote proportional rates of change in fecundity and mortality, respectively. Proportional rates are used because mortality and fecundity are measured on different scales. Senescence occurs when the proportional rate of change in mortality exceeds the proportional rate of change in fecundity. Non-senescence is subdivided into negative senescence, a strict subset in which the proportional rate of change in mortality is negative while the proportional rate of change in fecundity is stable or increasing, and negligible senescence, which encompasses all remaining cases of non-senescence.

Empirical evidence for negligible and negative senescence is taxonomically broad. Stable or declining adult mortality has been documented in long-lived bivalves such as the ocean quahog *Arctica islandica* (Ridgway and Richardson, 2011), colonial corals (Babcock, 1991), slow-growing elasmobranchs including the Greenland shark (Nielsen et al., 2016), and numerous plant and invertebrate species (Jones et al., 2014; Roper et al., 2021). Many of the species above show strictly negative actuarial senescence (mortality declines) while fecundity is stable or increases with age, so they meet the criterion for negative senescence (Babcock, 1991; Jones et al., 2014). A common feature across these taxa is indeterminate growth. Individuals continue to increase in body size after sexual maturity, and vital rates improve with size.

Life history models on ageing, often known as the disposable soma theory of ageing (Kirkwood, 1977; Cichon and Kozlowski, 2000; Kaplan and Robson, 2009; McNamara et al., 2009), provide a useful way to characterise how growth patterns co-evolve with mortality and fertility patterns. Most of these models, however, have focused on determinate growers. The few theoretical models that study indeterminate growth and negative senescence (Vaupel et al., 2004; Baudisch, 2008) rely on the growth–reproduction trade-off, in which body size and somatic condition are conflated, and somatic maintenance is absent as an allocation decision. Whether negative senescence remains an evolutionary outcome when maintenance competes directly with growth and reproduction has not been addressed, and the factors promoting it therefore remain unresolved.

Here we fill this gap. We formulate a three-dimensional optimal control problem in which an individual allocates energy among reproduction, somatic growth, and somatic maintenance. We assume that the soma accumulates damage over time, and investing in maintenance slows this process. We let mortality depend on both size and accumulated somatic damage. Using invasion analysis and Pontryagin’s maximum principle, we derive the uninvadable allocation strategy and a condition for negative actuarial senescence, and solve the resulting system numerically via GPOPS-II (Patterson and Rao, 2014). We show that indeterminate growth and negative senescence can arise as part of an uninvadable strategy, and we identify the conditions on allocation returns that produce this outcome. Section 2 formulates the model and optimality conditions. Section 3 presents the general conditions under which indeterminate growth and negative senescence emerge as uninvadable strategies, illustrated with representative numerical cases. Section 4 discusses the implications for the relationship between indeterminate growth, somatic maintenance, and senescence.

## 2 Model

### 2.1 Biological scenario

We consider a large panmictic population of haploid individuals reproducing asexually, with population size regulated by density-dependent competition affecting fecundity. The population is structured into continuous age classes, with individuals undergoing birth, development, reproduction, and death. Each individual at age *t* ∈ *T* = [0, ∞) is characterized by a two-dimensional physiological state **x**(*t*) = (*x*(*t*), d(*t*)) ∈ ℝ^2^, where *x*(*t*) represents somatic capital (e.g. body size, brain tissue, etc), which we will refer to as *size* from now on (see Table 1 for the list of symbols for the key variables and parameters of the model). The second state variable, d(*t*), describes the physiological condition of the soma, which we refer to as accumulated *damage* to the soma. Damage captures how the somatic condition deteriorates through processes such as oxidative stress, DNA damage, and cellular senescence, and we interpret d(*t*) as the state variable underlying physiological ageing, as it reflects mortality arising from imperfect somatic maintenance. Individuals acquire energy at a rate *P* (*x*(*t*)) = *a x*(*t*)^*c*^, which scales allometrically with size, reflecting Kleiber’s law (e.g., Kleiber, 1932; West et al., 1997) with *a >* 0 and *c <* 1. At each age *t*, individuals allocate their available energy among competing life history functions through a two-dimensional, age-dependent trait **u**(*t*) = (*u*_*b*_(*t*), *u*_*m*_(*t*)) ∈ [0, 1]^2^ : *u*_*b*_ + *u*_*m*_ ≤ 1, which we assume to be an evolving trait (control variable). Here *u*_*b*_(*t*) denotes the proportional allocation to reproduction and *u*_*m*_(*t*) denotes the proportional allocation to somatic maintenance. The remaining fraction 1 − *u*_*b*_(*t*) − *u*_*m*_(*t*) is allocated to growth. In Fig. 2, we have illustrated the life history model described here. We denote by **u** = *{***u**(*t*)*}*_*t*∈*T*_ ∈ *U* [*T*] the entire trait schedule expressed through an individual’s lifespan, where *U* [*T*] is a set of bounded and piecewise continuous functions on *T*. We follow standard invasion analysis for age-structured populations with infinite-dimensional traits (e.g. Day and Taylor, 2000; Metz et al., 2016; Avila et al., 2021; Avila and Lehmann, 2026; see also Appendix A).

**Table 1.** Variables and parameters of the model. Throughout, variables without subscript or superscript denote mutant values (e.g. **u**(*t*), **x**(*t*)); with superscript “*” (e.g. **u**^*\**^(*t*), **x**^*\**^(*t*)) denotes that the quantity is evaluated along the (candidate) uninvadable path; with subscript “r” denotes resident values (e.g. **u**_r_(*t*), **x**_r_(*t*))

| Symbol |  |
| --- | --- |
| $t$ | Age, $t \in \mathcal{T}$ |
| $\mathbf{u}(t)$ | Proportional allocation strategy $\mathbf{u}(t) = (u_b(t), u_m(t))$ (evolving trait). |
| $u_b(t)$ | Proportional allocation to reproduction. |
| $u_m(t)$ | Proportional allocation to somatic maintenance. |
| $\mathbf{x}(t)$ | Physiological state vector $\mathbf{x}(t) = (x(t), d(t))$. |
| $x(t)$ | Size or somatic capital. Growth rate $\dot{x}(t) = \alpha_x (1 - u_b(t) - u_m(t))^{\beta_x} P(x(t))$. |
| $d(t)$ | Somatic damage. Accumulation rate of somatic damage: $\dot{d}(t) = \alpha_d (\eta - u_m(t))^{\beta_d} P(x(t))$. |
| $l(t)$ | Survival probability to age $t$. Rate of change in survival: $\dot{l}(t) = -\mu(t) l(t)$. |
| $P(x)$ | Size-dependent production function $P(x) = ax^c$. |
| $b(t)$ | Age-specific fecundity rate $b(t) = \alpha_b u_b(t)^{\beta_b} P(x(t))$. |
| $\mu(t)$ | Age-specific mortality rate $\mu(\mathbf{x}(t)) = d(t) + \mu_x x(t)^\kappa + \mu_e$ |
| $\boldsymbol{\lambda}(t)$ | Costate vector $\boldsymbol{\lambda}(t) = (\lambda_x(t), \lambda_d(t), v(t))$ (here, always evaluated at the candidate uninvadable path). |
| $\lambda_x(t)$ | Costate variable associated with size; marginal fitness value of growth. |
| $\lambda_d(t)$ | Costate variable associated with somatic damage; marginal fitness cost of damage accumulation (negative). |
| $v(t)$ | Costate variable associated with survival probability; Fisher's reproductive value. |
| $f_c^*(t)$ | Fitness returns from current investments into reproduction and soma; $f_c^*(t) = b^*(t) + \dot{x}^*(t) \lambda_x^c(t) + \dot{d}^*(t) \lambda_d^c(t)$. |
| $\alpha_b, \alpha_x, \alpha_d$ | Conversion efficiency parameter for allocation to reproduction, growth and maintenance. |
| $\beta_b, \beta_x, \beta_d$ | Scaling factor for allocation to reproductive, growth and maintenance. |
| $a$ | Conversion efficiency parameter for size-dependent production. |
| $c$ | Scaling parameter of production ($c < 1$ implies diminishing returns). |
| $\eta$ | Proportion of resources needed to be allocated into maintenance in order to prevent any further somatic damage from accumulating. |
| $\kappa$ | Direction of size dependence in intrinsic mortality ($\kappa = -1$: larger size reduces mortality). |
| $\mu_e$ | Size-independent extrinsic mortality component. |
| $\mu_x$ | Scaling parameter for size-dependent intrinsic mortality. |

**Figure 2.**
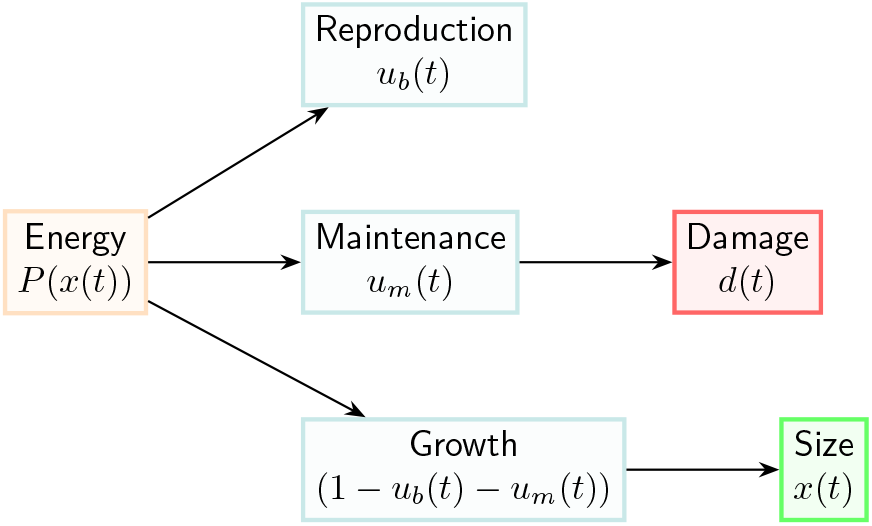
Schematic for the life history model. Energy production *P* (*x*(*t*)) scales allometrically with body size and is partitioned among three competing functions through proportional allocation traits to reproduction *u*_*b*_(*t*) and somatic maintenance *u*_*m*_(*t*). The residual fraction (1 − *u*_*b*_(*t*) − *u*_*m*_(*t*)) is allocated to growth, which increases size *x*(*t*). Maintenance allocation reduces the somatic damage *d*(*t*).

### 2.2 Invasion analysis

In order to characterise the invasion process, from now on, let **u** = *{***u**(*t*)*}*_*t*∈*T*_ and **x** = *{***x**(*t*)*}*_*t*∈*T*_ denote the mutant trait and state schedules, respectively, and let **u**_r_ = *{***u**_r_(*t*)*}*_*t*∈*T*_ and **x**_r_ = *{***x**_r_(*t*)*}*_*t*∈*T*_ denote the corresponding resident trait and state schedules. We consider the fate of a rare mutant individual with schedules (**u, x**) in a population monomorphic for resident individuals with schedules (**u**_r_, **x**_r_). We assume that the resident population is at demographic equilibrium and adopt standard invasion analysis assumptions, including a sufficiently large population size such that genetic drift can be neglected. From standard results on age-dependent branching processes (Crump and Mode, 1968, Corollary 5.1; Mode, 1971, Example 8.2), a mutant lineage descending from an ancestor with schedule **u** goes extinct in a monomorphic resident population with schedule **u**_r_ if and only if

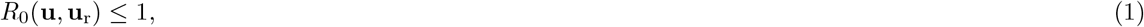

where *R*_0_(**u, u**_r_) is the basic reproductive number of a mutant individual. Namely, the expected number of surviving offspring produced by a mutant individual with trait schedule **u** in a resident population with trait schedule **u**_r_. Here, we assume that the basic reproductive number of the mutant takes the following form

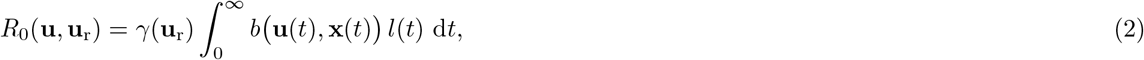

where *b* (**u**(*t*), **x**(*t*)) = *f* (*u*_*b*_(*t*)) *P* (*x*(*t*)) is the birth rate of a mutant individual, where *f* (*u*_*b*_) describes how reproductive allocation translates into fecundity and *P* (*x*(*t*)) is the size-dependent energy budget. In eq. (2) *γ*(**u**_r_) captures density-dependent regulation affecting fecundity, and *l*(*t*) denotes survival probability to age *t*, which satisfies

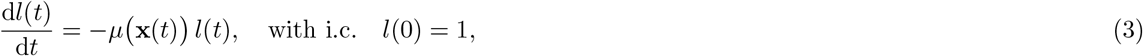

where “i.c.” stands for initial condition and the age-specific mortality rate *µ* (**x**(*t*)) is given by

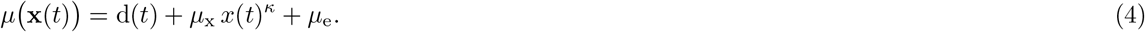

Here the first term represents the intrinsic mortality component arising from somatic damage d(*t*), while the second term represents the size-dependent intrinsic mortality component, with *κ* = −1 reflecting reduced mortality at larger sizes and *µ*_x_ *>* 0 scaling its contribution. Constant extrinsic mortality is denoted by *µ*_e_. The physiological state variables **x**(*t*) = (*x*(*t*), d(*t*)) evolve according to

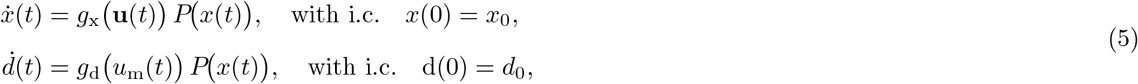

where 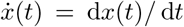 hereafter denotes the total derivative with respect to age *t* and *g*_x_(**u**(*t*)) and *g*_d_(*u*_m_(*t*)) describe how proportional allocation traits affect the corresponding state equations, with both scaled by the energy budget *P* (*x*(*t*)). Collectively, we refer to the survival and physiological state variables as the *state variables*, denoting them by **y** = (*l*(*t*), *x*(*t*), d(*t*)).

### 2.3 Uninvadability

A trait schedule **u**^*\**^ = *{***u**^*\**^(*t*)*}*_*t*∈*T*_ with an associated state schedule **y**^*\**^ = *{***y**^*\**^(*t*)*}*_*t*∈*T*_ is considered to be uninvadable if it cannot be displaced by any rare mutant; that is, if it solves the following maximisation problem

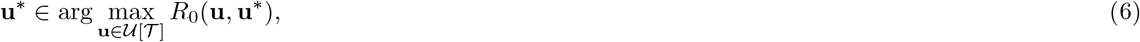

where the basic reproductive number of a mutant individual is given by eq. (2) and the maximisation is subject to the mutant dynamics constraints given by eqs. (3) and (5). We know from applications of optimal control theory (Bryson and Ho, 1975; Caputo, 2005; Weber, 2011; Kamien and Schwartz, 2012) to life history theory (e.g. Day and Taylor, 2000; Metz et al., 2016; Avila et al., 2021; Avila and Lehmann, 2026) that the maximisation problem above can be reformulated using Pontryagin’s Maximum Principle (see Appendix A and B) into maximisation of the Hamiltonian *H*(**u**(*t*), **x**^*\**^(*t*), ***λ***(*t*)) with respect to **u**(*t*) at each age *t* ∈ *T*, where Hamiltonian function

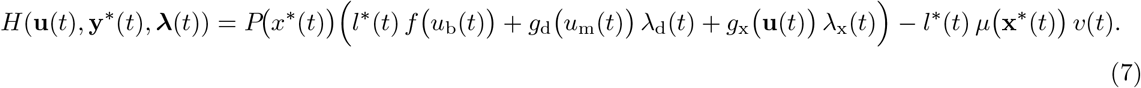

Here ***λ***(*t*) = (*λ*_x_(*t*), *λ*_d_(*t*), *v*(*t*)) is a vector of *costate variables* (or shadow values) associated with state variables **y**^*\**^ = (*x*^*\**^(*t*), d^*\**^(*t*), *l*^*\**^(*t*)), respectively, evaluated at the candidate uninvadable path. Each costate variable gives the fitness effect of marginally increasing the associated state variable (e.g. Dorfman, 1969, Perrin et al., 1993; Avila et al., 2021). The Hamiltonian decomposes fitness contributions from all life history activities: the first term captures immediate gains from reproduction, while the remaining terms capture future fitness consequences weighted by the costate variables: the second from somatic maintenance weighted by *λ*_d_(*t*), the third from growth weighted by *λ*_x_(*t*), and the fourth from survival weighted by *v*(*t*). The costate variables *λ*_d_(*t*), *λ*_x_(*t*), and *v*(*t*) represent, respectively, the marginal value of damage accumulation (always negative), the marginal value of growth, and Fisher’s (1930) *reproductive value* (i.e. the marginal value of surviving). Note that in the literature on life history theory with transfers, the concept of reproductive value has been extended to account for the fitness value of survival through future transfers and is known as the *value of life* (e.g., Kaplan and Robson, 2009). Note that in defining the Hamiltonian in eq. (7), we have factored out the density-dependent factor *γ*(**u**_r_) *>* 0 as it appears in *R*_0_(**u, u**_r_) as a positive multiplicative constant (recall (2)) and thus it does not affect the maximisation (6).

Here the costate dynamics evaluated at the uninvadable strategy are

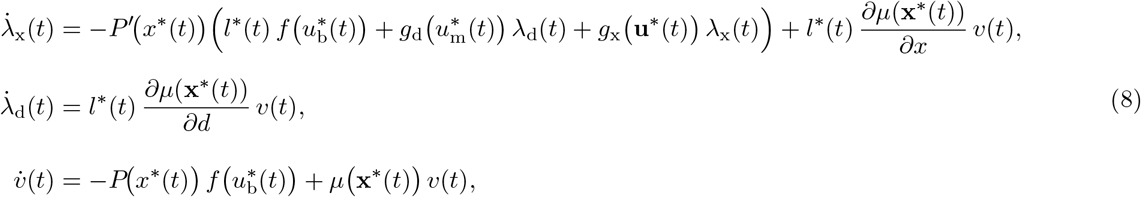

where *f* ^*′*^(*x*) ≡ d*f* (*x*)*/* d*x* notation throughout means a derivative with respect to its argument and lim_*t*→∞_ ***λ***(*t*) = 0.

#### 2.3.1 Uninvadable mortality rate and the condition for actuarial senescence

The uninvadable mortality rate *µ*^*\**^(*t*) = *µ*(**x**^*\**^(*t*)) (Avila and Lehmann, 2026; see Appendix A.1 for details) can be expressed as

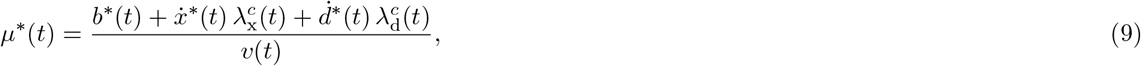

where we have suppressed writing the dependence on other variables besides age *t*, and 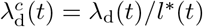 and 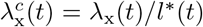 are current-value costates (costates conditional on survival, see e.g. Avila et al., 2021). Eq. (9) says that mortality varies inversely with the reproductive value and increases with the current reproduction, and returns from investment into current growth and maintenance. This result reflects the classic observation of Fisher (1930) that mortality tends to vary inversely with reproductive value, but (9) emphasises how fitness returns from current investments into reproduction also matter.

Let us define 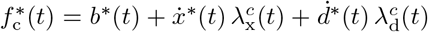 as the fitness returns from current investments into reproduction and soma. We show in Appendix A.2 that the uninvadable mortality rate increases when

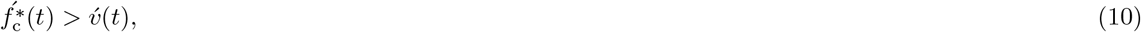

i.e. the proportional rate of change in fitness returns from current investments into reproduction and soma 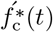 is higher than the proportional rate of change in the reproductive value 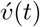. Subsituting eq. (9) in the dynamics for *v*(*t*) (eq. (8)) gives 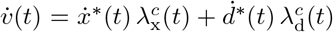. Thus, reproductive value increases when the fitness gain from growth exceeds the fitness cost of accumulating damage. This implies that reproductive value can continue to rise after maturity only under indeterminate growth; for determinate growers, once growth stops, 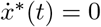 and *v*(*t*) declines from that age onwards. Thus, negative actuarial senescence can emerge when continued growth results in increasing reproductive value (future fitness potential), such that it outpaces the fitness returns from current investments into reproduction and soma 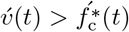.

### 2.4 Selection pressures

The age-specific selection gradients give the direction of selection for trait components *u*_*b*_(*t*) and *u*_*m*_(*t*)

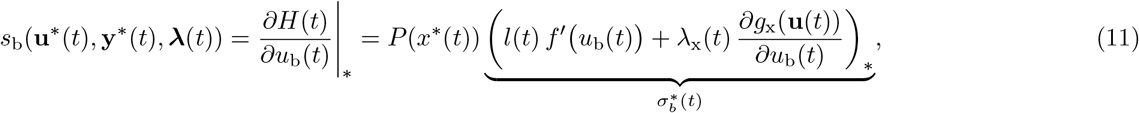

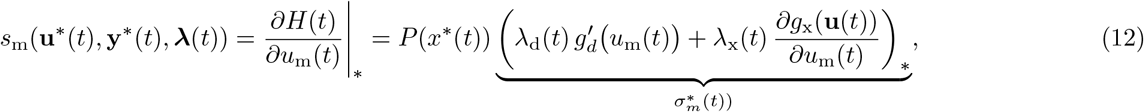

The selection gradients *s*_b_(*t*) and *s*_m_(*t*) describe the fitness consequences of reallocating resources at age *t* toward reproduction and somatic maintenance, respectively. For reproduction, the first term (inside parentheses) captures the direct fitness gain from offspring production, while for maintenance, it captures the future fitness gain from reduced damage accumulation weighted by *λ*_d_(*t*). In both cases, the second term represents the fitness cost of directing resources away from growth due to reallocation into reproduction or maintenance. Selection on allocation at age *t* thus reflects a trade-off between immediate or delayed fitness benefits from reproduction or maintenance and the future fitness cost of reduced growth. Here we have denoted 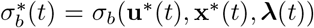 and 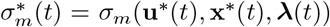 as the switching functions for reproduction and maintenance, respectively. This is because the sign of 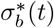 and 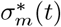 define the selection gradient as *P* (*x*^*\**^(*t*)) *>* 0.

### 2.5 Scaling of proportional allocation into life history functions

The allocation-response functions *f*(*u*_b_(*t*)), *g*_x_ (**u**(*t*)), and *g*_d_ (*u*_m_(*t*)) are phenomenological descriptions of how proportional allocation effort translates into the life history rates of fecundity, growth, and accumulation of damage, respectively. We treat these conversion processes as separable from energy supply, which scales allometrically according to *P* (*x*(*t*)), a standard assumption in life history models (e.g. Perrin, 1992; Perrin et al., 1993; Vaupel et al., 2004). We assume that the allocation-response functions follow a power law,

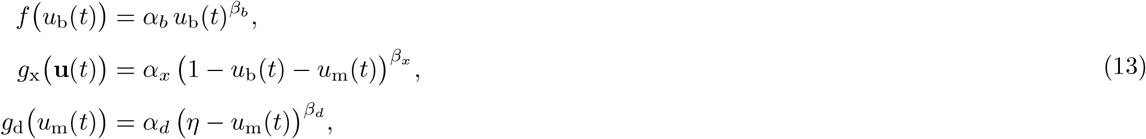

where *α*_*b*_, *α*_*x*_, *α*_*d*_ characterise the conversion efficiency of resource allocation effort into life history rates, and 0 *< η* ≤ 1 is the threshold giving the proportion of resources needed to be invested in maintenance in order to stop the damage accumulation. For our forthcoming analysis, we assume *η* = 1 in line with the classic disposable soma literature (see e.g. Cichon and Kozlowski, 2000, where this is also assumed), meaning that maintenance is costly and only full allocation would negate the damage completely. Here, *β*_*b*_, *β*_*x*_, *β*_*d*_ are scaling factors of the proportional allocation effort, and their effect on allocation-response functions is illustrated in Fig. 3.

**Figure 3.**
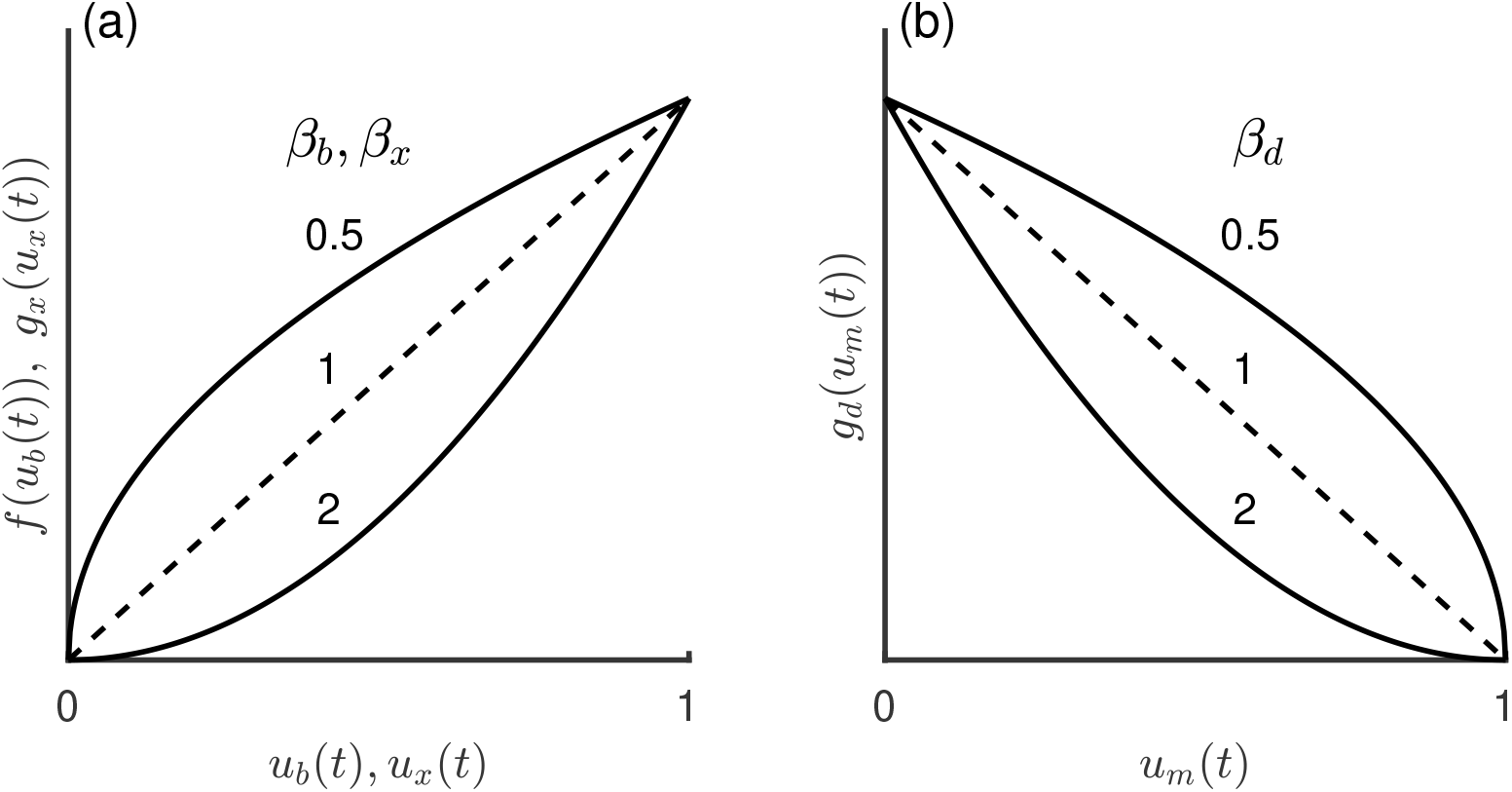
Scaling of the allocation-response functions for the three life history activities. Panel (a): the fecundity allocation-response function *f* (*u*_b_) and the growth allocation-response function *g*_x_(*u*_x_), both increasing in their respective allocation trait. Here, *u*_x_(*t*) = 1 − *u*_b_(*t*) − *u*_m_(*t*) the residual allocation to growth. Panel (b): the damage-accumulation allocation-response function. *g*_d_(*u*_m_), which decreases with maintenance allocation. The nonlinearity parameter *β* determines the shape of each function. For the increasing functions in (a), *β >* 1 corresponds to increasing marginal returns to allocation and *β <* 1 to diminishing marginal returns. For the decreasing function in (b) the interpretation is reversed: *β >* 1 implies diminishing marginal reductions in deterioration with increasing maintenance allocation, while *β <* 1 implies increasing marginal reductions.

In this formulation, allocation to somatic maintenance slows the accumulation of damage, which reduces intrinsic mortality arising from somatic damage d(*t*) at later ages. At the same time, allocating resources to maintenance reduces the energy available for reproduction and growth. Growth increases size *x*(*t*), which affects all three life history allocation functions through production but also modifies mortality through the size-dependent intrinsic mortality term.

## 3 Uninvadable life history outcomes

The life history model of Section 2, given by (2)–(5) and (13), produces a range of growth patterns (determinate versus indeterminate) and senescence patterns (negligible and negative senescence). In Appendix B, we characterise some general properties of the candidate uninvadable life history using Pontryagin’s maximum principle, and show how the scaling factors of resource allocation traits (*β*_*b*_, *β*_*x*_, *β*_*d*_) shape the uninvadable life history outcomes by determining the phases of the uninvadable life history schedule. Here we present the full numerical solutions obtained with the optimal control solver GPOPS-II (Patterson and Rao, 2014) and verify that they satisfy the analytical prediction set by the necessary condition for uninvadability (see the caption of Fig. 5 for details). We first examine two biologically relevant parameter combinations in detail, producing determinate growth with senescence (Section 3.1) and indeterminate growth with negative senescence (Section 3.2). We then present results from exploring uninvadable life-history outcomes across a broader range of combinations of scaling factors (*β*_*b*_, *β*_*x*_, *β*_*d*_) that are of biological significance (Section 3.3, see also Appendix C for further detail). This analysis allows us to highlight different pathways through which indeterminate growth and negative senescence can occur.

The first worked-out case presented in Section 3.1 assumes linear returns to reproduction and growth, *β*_*b*_ = *β*_*x*_ = 1, with diminishing returns to scale in maintenance, *β*_*d*_ = 4). Linear returns to reproduction and growth are assumed in many classical life history models, which motivates the first case (Macevicz and Oster, 1976; Vincent and Pulliam, 1980; Schaffer, 1982; Kozlowski, 1992; Cichon and Kozlowski, 2000; Day and Taylor, 2000; Avila et al., 2019). The second case presented in Section 3.2 assumes diminishing returns in all three life history functions (*β*_*b*_ = *β*_*x*_ = 0.5, *β*_*d*_ = 2), since there are many reasons to expect diminishing returns to reproduction and growth instead, such as gamete competition, structural constraints, and physiological ceilings in output (e.g. see discussion in Heino and Kaitala, 1999; Schärer, 2009; Frank, 2013). In both cases, we assume diminishing returns to somatic maintenance (*β*_*d*_ *>* 1), following the standard assumption in life history models that incorporate maintenance (e.g. Cichon and Kozlowski, 2000; Kaplan and Robson, 2009) and the disposable soma tradition (Kirkwood, 1977, 2017). This assumption is well-justified by empirical evidence from cellular repair processes, in which the cost of reducing molecular error rates increases with the accuracy achieved (Galas and Branscomb, 1978; Kirkwood, 2012).

### 3.1 Determinate growth and senescence

Linear returns to growth and reproduction lead to the so-called bang-bang allocation between growth and reproduction, a single switch that directs all non-maintenance energy first to growth and then to reproduction, resulting in determinate growth, while allocation to maintenance has an interior solution. The resulting life history allocation schedule is shown in Fig. 4 panel (a), with the associated growth and mortality trajectories in panel (b). It follows from eq. (11) and (13), assuming *β*_*b*_ = *β*_*x*_ = 1, that the sign of 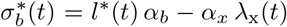 determines the growth and reproduction phases, where 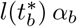 is the marginal benefit of reproduction and 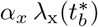 the marginal value of continued growth. Early in life 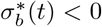, because the marginal value of growth is high (high *λ*_x_(*t*)). As the organism grows, *λ*_x_(*t*) falls until, at age 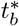, the gradient changes sign. The switch from growth to reproduction occurs where

**Figure 4.**
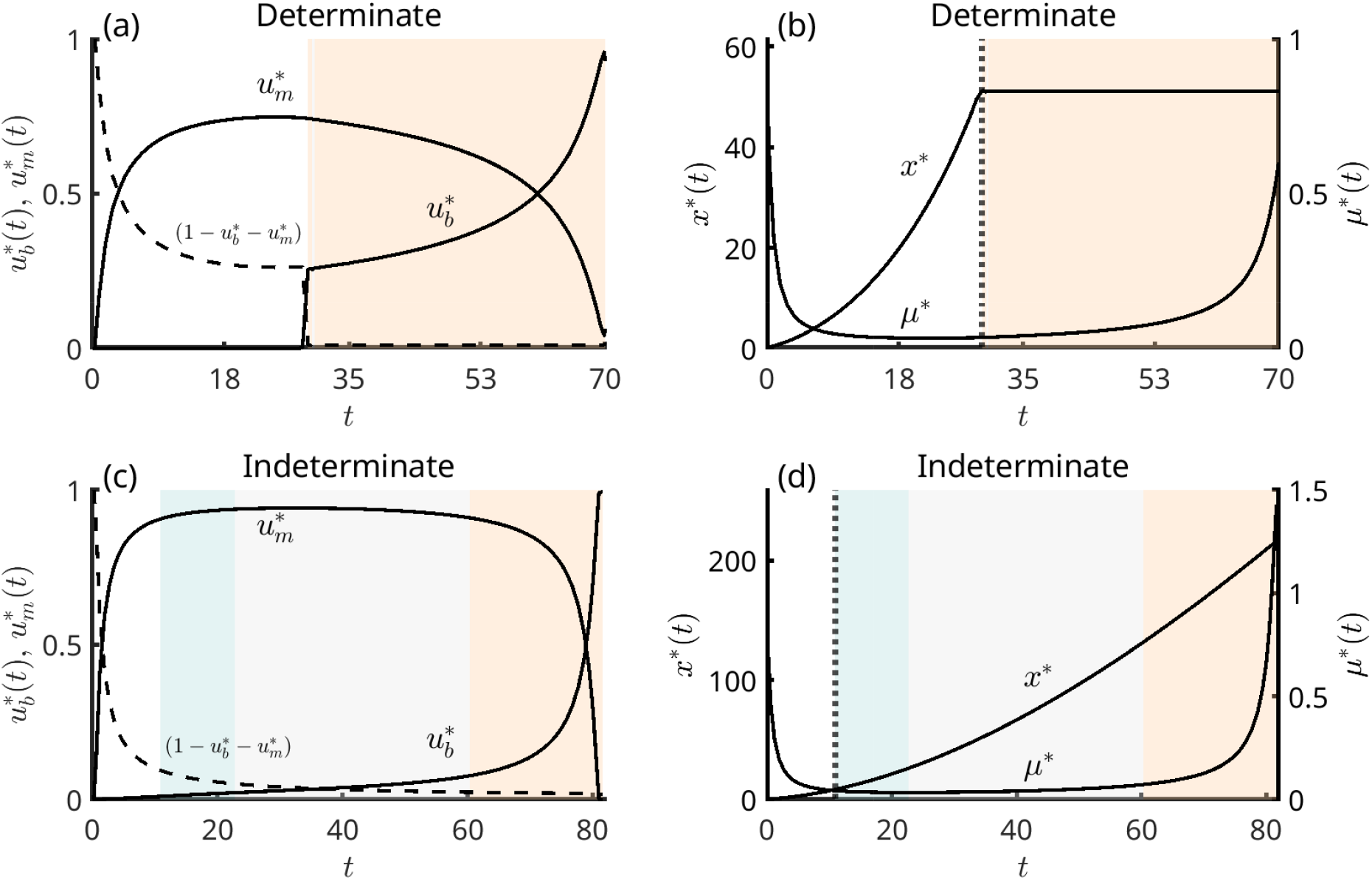
Two uninvadable life history outcomes. Panels (a) and (c) show energy allocation schedules; panels (b) and (d) show the corresponding size *x*^*\**^(*t*) and mortality *µ*^*\**^(*t*) trajectories. Background shading indicates the senescence phase: blue for negative senescence, grey for negligible senescence, and orange for senescence. (a)–(b) Determinate growth with senescence (*β*_*b*_ = *β*_*x*_ = 1, *β*_*d*_ = 4, *η* = 1): linear returns to reproduction and growth produce bang-bang allocation, with a switch from full growth allocation to full reproductive allocation at maturity. (c)–(d) Indeterminate growth with negative senescence (*β*_*b*_ = *β*_*x*_ = 0.5, *β*_*d*_ = 2, *η* = 1): diminishing returns to reproduction and growth sustain simultaneous allocation to growth and reproduction after maturity, producing a period of negative senescence followed by negligible senescence. Parameter values common to all cases: *a* = 1, *c* = 0.75, *α*_*b*_ = 0.35, *α*_*x*_ = 0.8, *µ*_e_ = 0.001, *µ*_x_ = 0.17, *κ* = − 1, *d*_0_ = 0.01, *x*_0_ = 0.2. Numerical solutions obtained using GPOPS-II with the hp-Patterson–Rao collocation method, IPOPT solver, mesh tolerance 10^−5^, and between 3 and 18 collocation points per mesh interval over a time horizon *t*_*f*_ = 1000.

**Figure 5.**
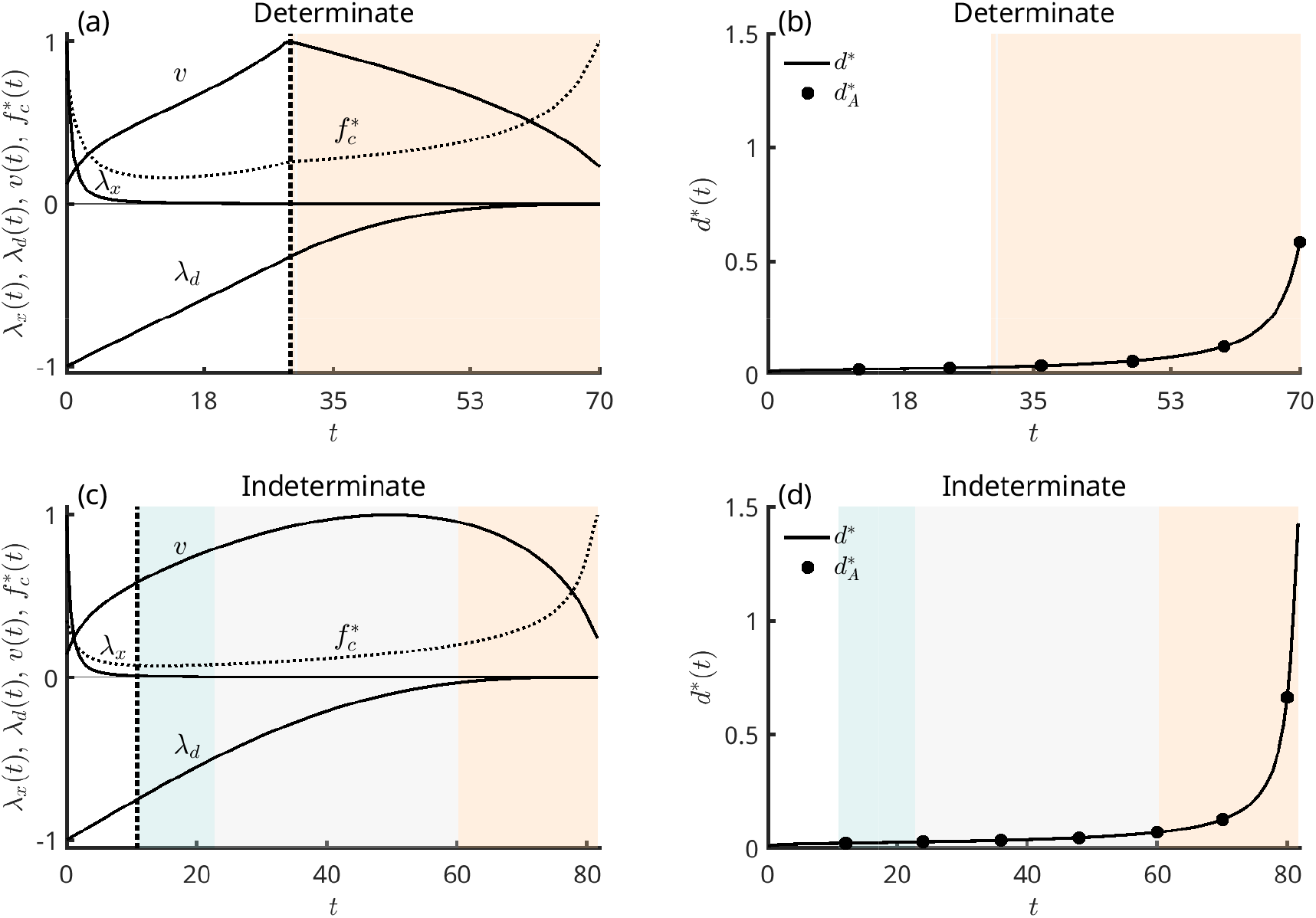
Costate values (shadow prices) associated with future growth *λ*_x_, accumulation of somatic damage *λ*_d_ and survival *v* for (a) determinate growth and (c) indeterminate growth. Background shading indicates the senescence phase: blue for negative senescence, grey for negligible senescence, and orange for senescence. Each costate is shown on a normalised scale from − 1 to 1, together with the fitness returns from current investments into reproduction and soma, 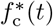 (dotted line). In the determinate case (a), the reproductive value *v* peaks at the age of maturity (dotted vertical line). In contrast, under indeterminate growth (c), *v* continues to increase after maturity, coinciding with the period of negative senescence as predicted in results presented in Section 2.3.1. Panels (b) and (d) show somatic damage *d*^*\**^(*t*) on the uninvadable path for the corresponding cases. The solid line *d*^*\**^ is obtained by GPOPS-II. Dots show 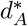 derived analytically from the condition *H*^*\**^ = 0. Since the two trajectories align in panels (b) and (d), indicating that the numerical solution for the entire schedule of (**u**^*\**^, **x**^*\**^) is consistent with analytical predictions. Parameter values as in Fig. 4.

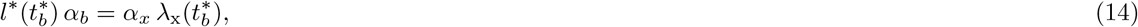

which defines the age at maturity 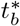. Singular arcs, in which eq. (14) holds over an interval rather than at an isolated point, cannot be formally excluded, but we do not consider them, as we did not observe them numerically; such knife-edge solutions arise only under special conditions (see Perrin and Sibly, 1993). While growth and reproduction allocation are bang-bang, maintenance takes an interior value, obtained by solving 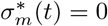 (eq. (12)). The closed-form solution is complicated in this case, but we observe that investment into maintenance is a concave function (Fig. 4 panel (a)). Fig. 5 panel (b) shows that the somatic damage *d*^*\**^(*t*) increases monotonically through life and accelerates once maintenance allocation falls to zero and reproduction increases (Fig. 4 panel (a)).

The corresponding uninvadable mortality *µ*^*\**^(*t*) is U-shaped (Fig. 4 panel (b)) and varies inversely with the reproductive value *v*(*t*) (Fig. 5 panel (a)), as predicted analytically by eq. (10). The fitness returns from current investments into reproduction and soma, 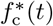, also follow a U-shaped pattern over the same period (Fig. 5 panel (a)). Early in life they are dominated by the growth term 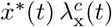, which declines as 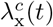 falls faster than growth itself slows. Later in life they are dominated by the rising reproduction term *b*^*\**^(*t*), once allocation switches to reproduction at 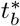. This switch in which term dominates produces the small change in slope visible in 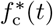 at 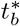. Mortality rate 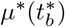 is lowest when 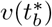 peaks, which is at reproductive maturity 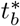. From Fig. 5 panel (a), the marginal value of growth *λ*_x_(*t*) and the marginal cost of damage accumulation *λ*_*d*_(*t*) both decline in magnitude with age. We verify the numerical solution against an analytical prediction. We do this by solving *H*^*\**^(*t*) = 0 (see Appendix A for details about Pontryagin’s maximum principle) for the somatic damage, which yields a closed-form trajectory 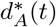 that coincides with the GPOPS-II output (see bullets in Fig. 5 panel (b)), confirming that the numerical and analytical predictions agree.

In summary, when reproduction and growth scale linearly with allocation (*β*_*b*_ = *β*_*x*_ = 1) while maintenance has diminishing returns (*β*_*d*_ = 4), the uninvadable life history recovers the classic pattern of growth followed by reproduction. Mortality is U-shaped, reaching its minimum at reproductive maturity 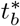, while the somatic damage *d*^*\**^(*t*) increases monotonically, rising rapidly late in life as maintenance allocation falls to zero and reproduction reaches its maximum. These results are consistent with classical disposable soma theory, and here we obtain qualitatively similar results to the model of e.g. Cichon and Kozlowski (2000).

### 3.2 Indeterminate growth and negative senescence

Diminishing returns from energy to growth and reproduction give rise to an indeterminate growth schedule (both growth and reproduction have interior solutions). The resulting life history allocation schedule is shown in Fig. 4 panel (c), with the associated growth and mortality trajectories in Fig. 4 panel (d). Here, the uninvadable life history schedule can be characterised by solving 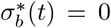 and 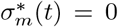 simultaneously for 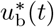 and 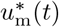 (recall eq. (11) and eq. (12)); but the closed-form solution is complicated in this case. In this scenario, selection favours indeterminate growth through two mechanisms. The first is diminishing returns to reproduction (*β*_*b*_ *<* 1): when the marginal fitness gain from each additional unit of reproductive investment decreases, going all-in on reproduction at maturity is wasteful, and it pays to keep allocating to growth. The second is diminishing returns to growth (*β*_*x*_ *<* 1): the marginal fitness cost of each unit withdrawn from growth increases as growth allocation gets smaller, so the organism continues to allocate to growth alongside reproduction rather than switching away from it entirely at maturity. The two mechanisms are complementary, both favouring continued growth after sexual maturity. Here, reproduction starts earlier than in the determinate case but stays at low levels for much of the life course, and growth is much slower (Fig. 4 panels (a) and (c)).

Similarly to the determinate case, allocation to maintenance is concave, but it stays much higher throughout most of the life course (Fig. 4 panels (a) and (c)). As in the determinate case, maintenance eventually drops rapidly as allocation to reproduction rises later in life. The resulting somatic damage *d*^*\**^(*t*) increases monotonically through life and accelerates once maintenance allocation falls to zero and reproduction increases (Fig. 5 panel (d)).

The corresponding uninvadable mortality *µ*^*\**^(*t*) is U-shaped but more rectangular (Fig. 4 panel (d)): it remains nearly constant after reproductive maturity and takes its lowest value before reproductive value peaks (Fig. 5 panel (c)). Similarly to the determinate case, 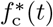 follows a U-shaped pattern (Fig. 5 panel (c)). As in the determinate case, mortality eventually increases with somatic damage, due to external mortality and relatively costly somatic maintenance (see Avila and Lehmann, 2026 for how lowering the maintenance cost can lead to steady-state life history). We observe a period of negative senescence after reproductive maturity, when mortality continues to decrease (blue region in Fig. 4). This result obtains because mortality decreases with size in our model (*κ* = −1): continued growth after maturity drives mortality down, while sustained maintenance keeps somatic damage from accumulating rapidly (Fig. 5 panel (d)). The net effect is declining mortality alongside increasing fecundity (Fig. 4 panel (d)). Similarly to the case with determinate growth, the marginal value of growth *λ*_x_(*t*) and the marginal cost of damage accumulation *λ*_*d*_(*t*) both decline in magnitude with age (Fig. 5 panel (c)).

In summary, when reproduction and growth have diminishing returns to allocation (*β*_*b*_ = *β*_*x*_ = 0.5) while maintenance also has diminishing returns (*β*_*d*_ = 2), the uninvadable life history recovers a pattern seen in many indeterminate growers that exhibit negative senescence (Jones et al., 2014). Mortality is U-shaped but more rectangular, reaching its minimum after reproductive maturity owing to continued investment in growth, while the somatic damage *d*^*\**^(*t*) increases slowly and rises rapidly only late in life as maintenance allocation falls to zero and reproduction reaches its maximum. These results extend classical disposable soma theory to allow negative senescence, driven by continued growth after maturity and by mortality scaling inversely with size.

### 3.3 Conditions for indeterminate growth and negative senescence

Beyond these two worked-out cases presented above, the scaling factors of resource allocation traits (*β*_*b*_, *β*_*x*_, *β*_*d*_) determine which uninvadable life history phases arise (see Appendix B). Varying them across the full set of linear and diminishing-returns combinations (Appendix C.1, Table 2) yields three general results. First, growth is determinate, i.e. a switch from pure growth to pure reproduction at maturity obtains only when both fecundity and growth scale linearly (*β*_*b*_ = *β*_*x*_ = 1). Diminishing returns to either reproduction or growth (*β*_*b*_ *<* 1 or *β*_*x*_ *<* 1) yield indeterminate growth, for two distinct reasons; namely, *β*_*b*_ *<* 1 penalises concentrating a high level of allocation on reproduction, whereas *β*_*x*_ *<* 1 penalises abandoning growth entirely. Second, positive maintenance appears on the uninvadable path only when *β*_*d*_ *>* 1, or when reproduction and growth both have diminishing returns (*β*_*b*_ *<* 1 and *β*_*x*_ *<* 1); in the latter case, high allocation to either growth or reproduction yields progressively smaller marginal returns, so at the margin, energy is better spent on maintenance, which is therefore selected at a positive interior value even though its own returns are only linear (Table 2g). Unlike Cichon and Kozlowski (2000), where growth and reproduction are always linear splits of production and only repair carries diminishing returns, our result shows that maintenance can be favoured at a positive level without diminishing returns to maintenance itself, provided both growth and reproduction have diminishing returns. Third, negative senescence arises in our model whenever continued growth lowers the size-dependent mortality term (*κ* = −1) faster than damage *d*(*t*) increases. The necessary condition for this is indeterminate growth (*β*_*b*_ *<* 1 or *β*_*x*_ *<* 1); early after reaching maturity, the decline in mortality can hold even without allocation to maintenance, since damage *d*(*t*) has not yet accumulated. Diminishing returns to reproduction (*β*_*b*_ *<* 1) extend this period by keeping growth allocation high after maturity. Because *β*_*b*_ *<* 1 already holds here, the general condition for positive maintenance given above reduces to *β*_*x*_ *<* 1 or *β*_*d*_ *>* 1, and whenever it is met, maintenance extends the period further by holding *d*(*t*) down. Lower extrinsic mortality *µ*_e_ extends the period still further by raising the fitness value of survival to old age. A period of negative senescence is eventually followed by senescence, because we assume that somatic maintenance is costly enough such that damage *d*(*t*) inevitably accumulates (in line with classic assumptions of disposable soma theory, see e.g. Cichon and Kozlowski, 2000).

**Table 2.** Uninvadable life histories for the eight parameter sets shown in Fig. 6. Parameters held constant across all runs: *a* = 1, *c* = 0.75, *α*_*d*_ = 0.01, *µ*_e_ = 0.001, *κ* = −1, *d*_0_ = 0.01, *x*_0_ = 0.2. The value of *α*_*b*_ is selected for each run so that the numerical objective *R*_0_ = 1 at the uninvadable solution. As *α*_*b*_ only scales the objective function, this choice does not affect the qualitative life history outcome. Life history phases are defined by which allocations are positive on the uninvadable path: Phase I (growth only), Phase II (growth and maintenance), Phase III (growth, reproduction, and maintenance), Phase IV (growth and reproduction), Phase V (reproduction and maintenance), Phase VI (reproduction only). NonS: negligible senescence. NegS: negative senescence. Cases marked (✓) satisfy the non-senescence condition only briefly after the onset of reproduction. Growth temporarily reduces mortality through the size-dependent term, but this decline is short-lived and does not constitute a sustained period of non-senescence.

| | $\beta_b$ | $\beta_x$ | $\beta_d$ | Life history phases | | Other parameters | Senescence | |
| --- | --- | --- | --- | --- | --- | --- | --- | --- |
|  |  |  |  | Pre-maturity | Post-maturity |  | NonS | NegS |
| (a) | 1 | 1 | 1 | I | VI | $\alpha_b = 0.96, \alpha_x = 0.30, \mu_x = 0.15$ | | |
| (b) | 1 | 1 | 4 | I → II | V → VI | $\alpha_b = 0.28, \alpha_x = 0.47, \mu_x = 0.15$ | | |
| (c) | 1 | 0.5 | 1 | I | IV → VI | $\alpha_b = 0.56, \alpha_x = 0.25, \mu_x = 0.10$ | (✓) | (✓) |
| (d) | 0.75 | 1 | 1 | I | IV → VI | $\alpha_b = 0.34, \alpha_x = 0.30, \mu_x = 0.07$ | (✓) | (✓) |
| (e) | 1 | 0.5 | 3 | II | III → VI | $\alpha_b = 0.48, \alpha_x = 0.35, \mu_x = 0.15$ | ✓ | (✓) |
| (f) | 0.5 | 1 | 3 | II | III → V → VI | $\alpha_b = 0.48, \alpha_x = 0.30, \mu_x = 0.10$ | ✓ | ✓ |
| (g) | 0.25 | 0.5 | 1 | II | III → V | $\alpha_b = 0.48, \alpha_x = 2.40, \mu_x = 0.18$ | ✓ | ✓ |
| (h) | 0.5 | 0.5 | 2 | II | III → IV → VI | $\alpha_b = 0.62, \alpha_x = 0.45, \mu_x = 0.18$ | ✓ | ✓ |

## 4 Discussion

We formulated a life-history model in which an individual allocates energy among reproduction, somatic growth, and somatic maintenance, with mortality decreasing with body size and increasing with somatic damage. Using Pontryagin’s maximum principle together with invasion analysis, we derived the necessary conditions for an uninvadable allocation strategy, an explicit expression for uninvadable mortality schedule (eq. 9), and a condition for actuarial senescence (eq. 10). Our analysis showed that scaling factors of resource allocation traits play a crucial role in determining the uninvadable life history schedules, and we solved the resulting system numerically for different biologically relevant parametrisations of these scaling factors.

The maximum principle yields an explicit expression for the uninvadable mortality rate: the fitness returns from current investment in reproduction and soma, divided by reproductive value (eq. 9). Mortality therefore varies inversely with reproductive value and rises with current returns, so that at each age it balances reproduction and somatic investment against survival. Differentiating this expression gives a condition for actuarial senescence (eq. 10): mortality increases with age whenever the proportional rate of change in fitness returns from reproduction and somatic investments exceeds that of reproductive value. In the special case where fitness returns from current investments in reproduction and soma remain roughly constant, mortality varies inversely with reproductive value, thereby recovering Fisher’s (1930) prediction. Negative (actuarial) senescence is the converse: mortality declines when reproductive value rises faster than fitness returns from current investments. In our model, this requires body size to keep increasing after maturity, because growth lowers size-dependent mortality (*κ* = −1) and raises future production, both of which increase reproductive value. The condition can thus be met only under indeterminate growth, which itself arises when returns to reproduction or to growth diminish with allocation effort.

Our numerical analysis shows that diminishing returns to reproduction (*β*_*b*_ *<* 1) and to growth (*β*_*x*_ *<* 1) can each promote negative senescence independently, and that the two can also act together. Both favour indeterminate growth, and because mortality is assumed to decrease with size (*κ* = −1), continued growth after maturity lowers mortality, producing a period of negative senescence. This extends previous work on two-dimensional growth–reproduction models, in which indeterminate growth arises from diminishing returns to reproduction (Sibly et al., 1985; Johansson et al., 2018; Vaupel et al., 2004) or arises when reproductive growth is restricted (Kozlowski and Ziólko, 1988), or from seasonality (Kozlowski, 1996). Of these, only Vaupel et al. (2004) connects indeterminate growth to negative senescence, but as it does not model maintenance allocation and somatic damage accumulation, it does not formally model the ageing process of a deteriorating soma. Vaupel et al. (2004) also only considers diminishing returns to reproduction. Our results thus extend the disposable soma theory (Kirkwood, 1977; Cichon and Kozlowski, 2000) to organisms with indeterminate growth and show that diminishing returns from both reproduction or growth can give rise to negative senescence. In our model, senescence nonetheless eventually sets in because, in line with the disposable soma theory, we assume maintenance is costly: all resources must be allocated to maintenance to prevent damage from accumulating (following e.g. Cichon and Kozlowski (2000) in this assumption). See Avila and Lehmann (2026) for a case where negligible senescence occurs under a determinate growth schedule when the cost of maintenance is low enough.

In our numerical analysis, we focused on diminishing returns of allocation traits rather than accelerating returns, since diminishing returns appear to be common in natural systems. However, accelerating returns to reproduction (*β*_*b*_ *>* 1) can be considered under some scenarios. For example, in some plants larger investment in reproduction might attract disproportionately more seed dispersers or pollinators, producing accelerating returns (Sallabanks, 1992; Schaffer and Schaffer, 1979). We show a numerical example of this in Appendix C.2 and find that when returns to reproduction accelerate, we obtain a determinate growth schedule and senescence.

In conclusion, our results, which extend the disposable soma theory to organisms with indeterminate growth and negative senescence, yield testable predictions. Species or populations in which returns to reproduction or to growth diminish strongly with allocation, and in which larger individuals have lower mortality, should show a stronger tendency toward negative or negligible senescence than those in which both returns are close to linear. Testing this requires longitudinal demographic and survival data (e.g. da Silva et al., 2022) alongside data on how reproductive allocation and growth rates change with size (e.g. Wenk and Falster, 2015). An interesting extension of our model would be to consider sexual reproduction and dimorphism in allocation between the sexes.

## Author Contributions

Arttu Soukainen: conceptualisation, methodology, formal analysis, numerical simulations, software, investigation, validation, writing – original draft, review & editing, visualisation. Piret Avila: conceptualisation, methodology, formal analysis, supervision, writing – original draft, review & editing, resources, funding.

## Data and resource availability

The MATLAB code implementing the optimal control solutions (GPOPS-II with ADiGator automatic differentiation), used to generate all numerical results and figures in this manuscript, will be made available upon publication.

## A Application of the Maximum principle for the life history problem

In this appendix, we apply Pontryagin’s maximum principle to the life history problem stated in the main text (eqs. (2)–(5)), which we restate here for readability. From optimal control theory (Bryson and Ho, 1975; Caputo, 2005; Weber, 2011; Kamien and Schwartz, 2012), and following its established use in life-history theory (León, 1976; Perrin, 1992; Day and Taylor, 2000; Metz et al., 2016; Avila et al., 2021), we derive the necessary conditions for an uninvadable life history. We follow Avila and Lehmann (2026) to derive the mortality rate on the uninvadable path. We then derive the condition under which actuarial senescence occurs.

Recalling eq. (2) of the main text, the fitness proxy here is given by the basic reproductive number, which can be written as

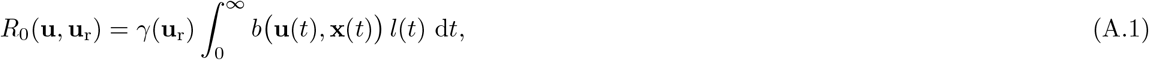

where *b* (**u**(*t*), **x**(*t*)) = *f* (*u*_*b*_(*t*)) *P* (*x*(*t*)). The basic reproductive number eq. (A.1) is subject to dynamic constraints of three state variables. First, the survival

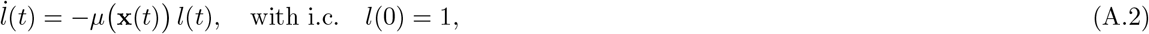

and physiological state variables **x**(*t*) = (*x*(*t*), d(*t*))

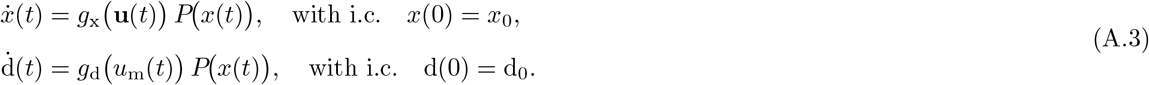

An uninvadable life history schedule **u**^*\**^ maximises the basic reproductive number, i.e.

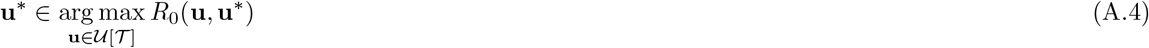

subject to the state dynamics in eqs. (A.2)–(A.3) evaluated along **u**^*\**^. Notice here that the fitness proxy is multiplicatively separable in mutant and resident quantities 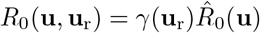, where

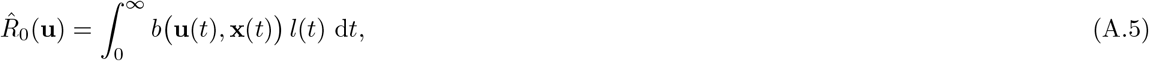

and thus maximising *R*_0_(**u, u**_r_) with respect to **u** is akin to maximising 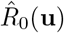 with respect to **u** (Metz et al., 2008). That is, solving for the uninvadable life history becomes a pure optimisation problem.

Now, supposing that the control schedule **u**^*\**^ with associated state schedule **y**^*\**^ is uninvadable. The maximum principle states that, then it is necessary that the control function **u**^*\**^(*t*) maximises the Hamiltonian for all *t* ∈ *T* :

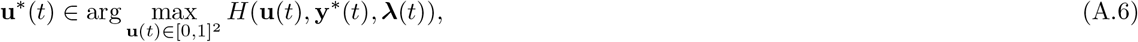

where the Hamiltonian for this optimisation problem takes the form

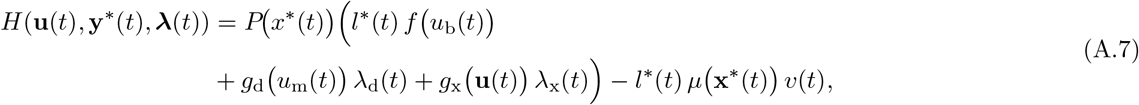

where the costate variables ***λ***(*t*) = (*λ*_x_(*t*), *λ*_d_(*t*), *v*(*t*)) are given by

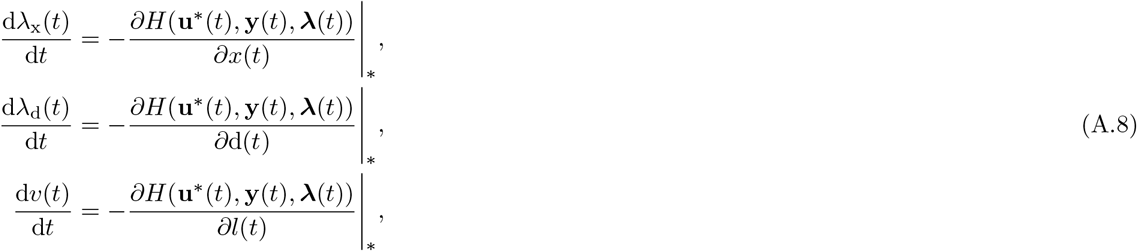

with transversality conditions *v*(0) = 1 and lim_*t*→∞_ *λ*_x_(*t*) = 0 and lim_*t*→∞_ *λ*_d_(*t*) = 0. eq. (A.6) yields the following first-order condition for the trait component *u*_*i*_(*t*) ∈ *{u*_b_(*t*), *u*_m_(*t*)*}* to be uninvadable

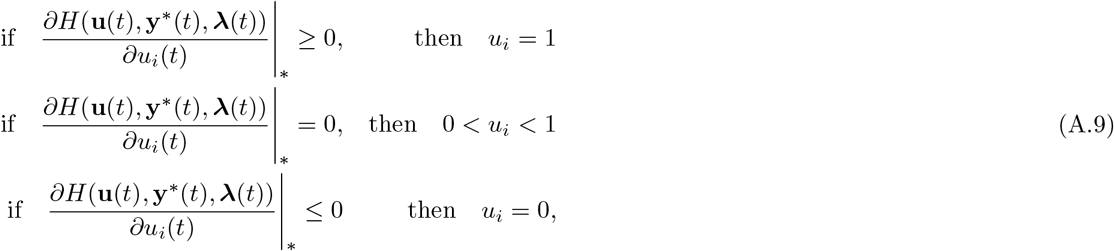

where the “|_*\**_” denotes that all variables are evaluated at the candidate uninvadable state. When the Hamiltonian is linear in the control *u*_*i*_(*t*) ∈ [0, 1], the inequalities become strict except at isolated points or at period intervals. In this case

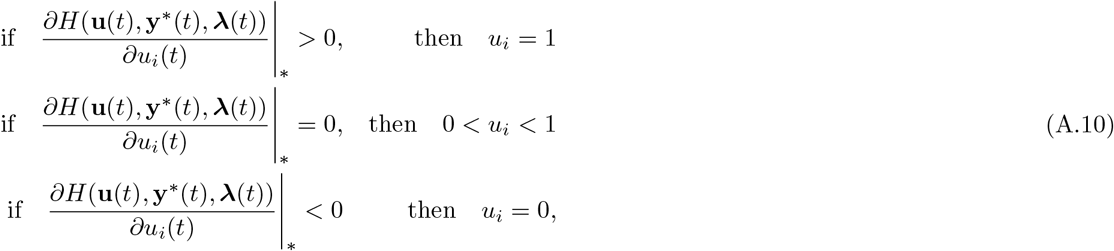

This yields a bang-bang solution in which the control switches between boundary values, except when the selection gradient vanishes. When the gradient remains zero over an interval (called a singular arc), the control takes an interior value 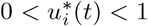. The singular control can be determined by repeatedly differentiating the condition *∂H*(*t*)*/∂u*_*i*_(*t*)|_*\**_ = 0 with respect to time

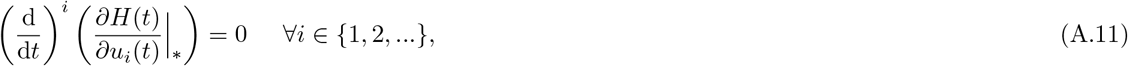

until the control *u*_*i*_(*t*) appears explicitly in the equation (e.g., Kopp and Moyer, 1965).

### A.1 Mortality on the uninvadable path

For autonomous infinite-horizon optimal control problems, the maximised Hamiltonian *H*^*\**^(*t*) ≡ *H*(**u**^*\**^(*t*), **y**^*\**^(*t*), ***λ***(*t*)) evaluated along the optimal trajectory vanishes identically (see Caputo (2005) for the general theory and Avila and Lehmann (2026) for applications in evolutionary biology). Thus, we have that

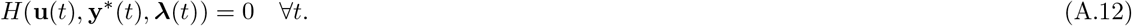

As demonstrated by Avila and Lehmann (2026) for a more general life history scenario, this constraint yields an explicit relationship for the optimal mortality rate

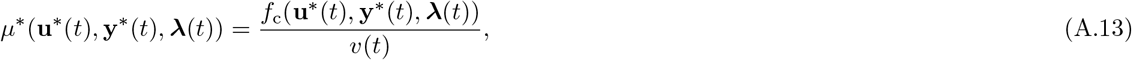

where

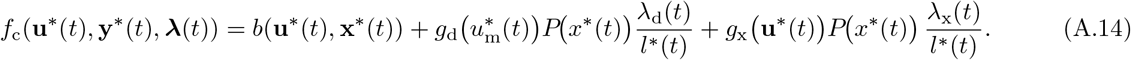

### A.2 Derivation of the senescence condition

We here derive the condition for senescence from the optimal mortality schedule eq. (A.13). To obtain a condition under which senescence occurs (i.e., d*µ*(*t*)*/* d*t >* 0), we take the total time derivative of eq. (A.13). Using shorthand notation 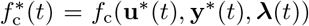 and *µ*^*\**^(*t*) = *µ*(**x**^*\**^(*t*)), we have from eq. (A.13) that

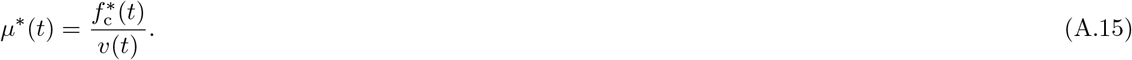

Applying the quotient rule yields

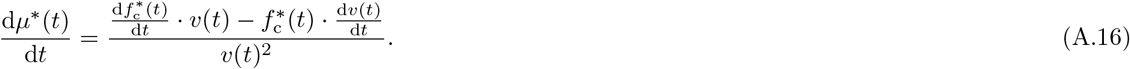

For senescence to occur, we require d*µ*^*\**^(*t*)*/* d*t >* 0, thus we have that

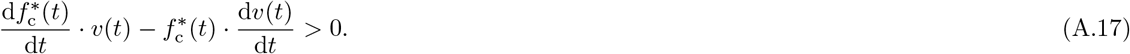

Rearranging and dividing both sides by *f*_c_(*t*) · *v*(*t*) *>* 0 yields

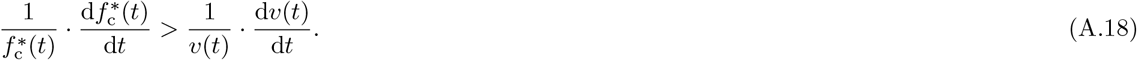

Using the notation of the main text that 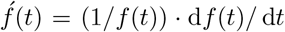 for any function *f* (*t*), we arrive at the final condition

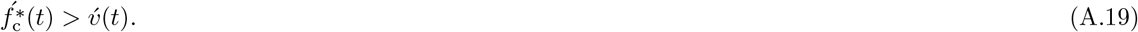

## B The role of scaling factors in determining the general properties of uninvadable life histories

In this Appendix, we derive some properties of the uninvadable life history schedule **u**^*\**^ for the model presented in the main text (eqs. 2–5). In particular, we focus on the role of the scaling factors of allocation (*β*_*b*_, *β*_*d*_, *β*_*x*_) in determining the possible phases of the uninvadable life history schedule. We do this by applying the necessary conditions for uninvadability obtained in Appendix A. Differentiating the Hamiltonian (7) with respect to **u**(*t*) and substituting the power-law allocation functions (13) yields the selection gradients

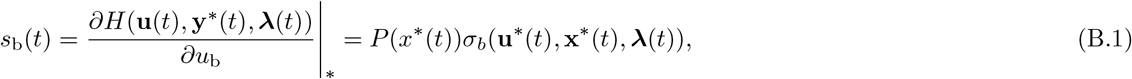

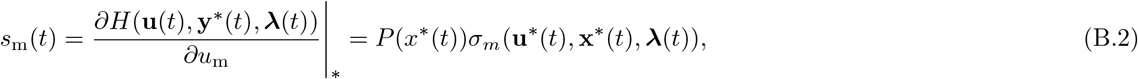

where

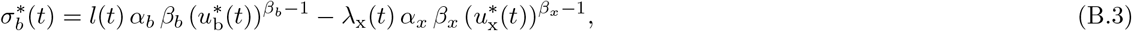

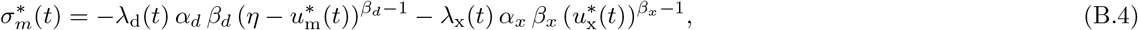

and 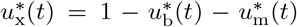 denotes the residual allocation to growth. Recall that *l*(*t*) *>* 0 and *λ*_x_(*t*) *>* 0 while *λ*_d_(*t*) *<* 0 throughout, so both switching functions balances a fitness gain against the fitness cost of reduced growth. Because *P* (*x*^*\**^(*t*)) *>* 0, the signs of the selection gradients *s*_b_(*t*) and *s*_m_(*t*) are determined entirely by the signs of the switching functions 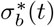 and 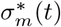. Applying the first-order conditions (A.9)–(A.10), each combination of signs of these switching functions corresponds to a distinct life history phase, defined by which allocations are positive on the uninvadable path:

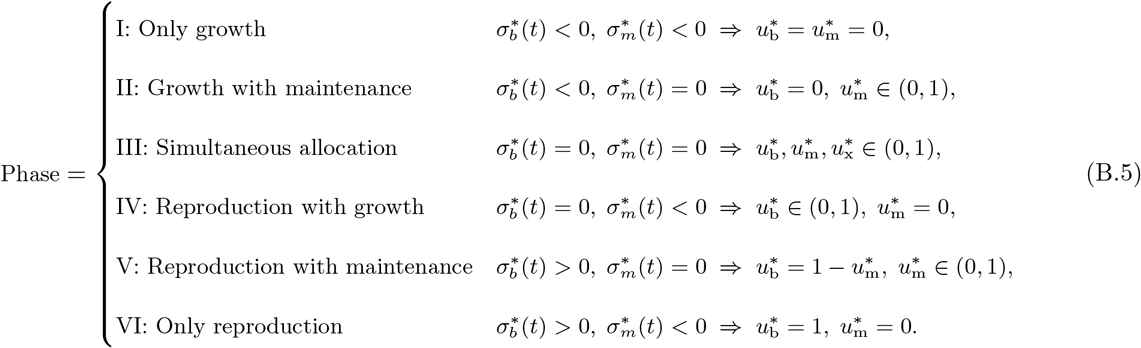

Which of these phases can arise on the uninvadable path depends critically on whether the controls enter the Hamiltonian linearly or nonlinearly. Standard results from optimal control theory (e.g., Bryson and Ho, 1975) establish that when a control variable enters the Hamiltonian linearly, the uninvadable solution lies on the boundary of the admissible set, except along singular arcs, which we do not consider here (as the singular arc solutions require many simultaneous constraints to hold and we did not observe them numerically). In our model, *u*_b_(*t*) enters linearly when *β*_*b*_ = *β*_*x*_ = 1, because both terms of 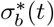 then become independent of 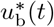; analogously, *u*_m_(*t*) enters linearly when *β*_*d*_ = *β*_*x*_ = 1. Under these conditions 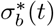 and 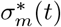 retain a definite sign that is independent of the controls, so only boundary allocations are uninvadable. When at least one of the relevant scaling factors departs from unity, the corresponding control enters nonlinearly, and interior solutions become possible. When this occurs for both controls, we can obtain a solution consisting of a time interval during which growth, reproduction, and maintenance are positive simultaneously. The scaling factors *β*_*b*_, *β*_*x*_, and *β*_*d*_ therefore play a crucial role in determining the life history phases along the uninvadable path.

## C Numerical analysis

### C.1 Diminishing and linear returns to scale

In this appendix we present selected examples from our numerical analysis representing all eight combinations of linear (*β* = 1) and diminishing (*β <* 1) returns across the three scaling factors (Fig. 6 and Table 2). Specific parameter values were selected to produce well-separated numerical examples to highlight the main properties of the uninvadable life history schedule **u**^*\**^ for each combination. These are independent runs and need not match the parameter values used in Figs. 4 and 5.

**Figure 6.**
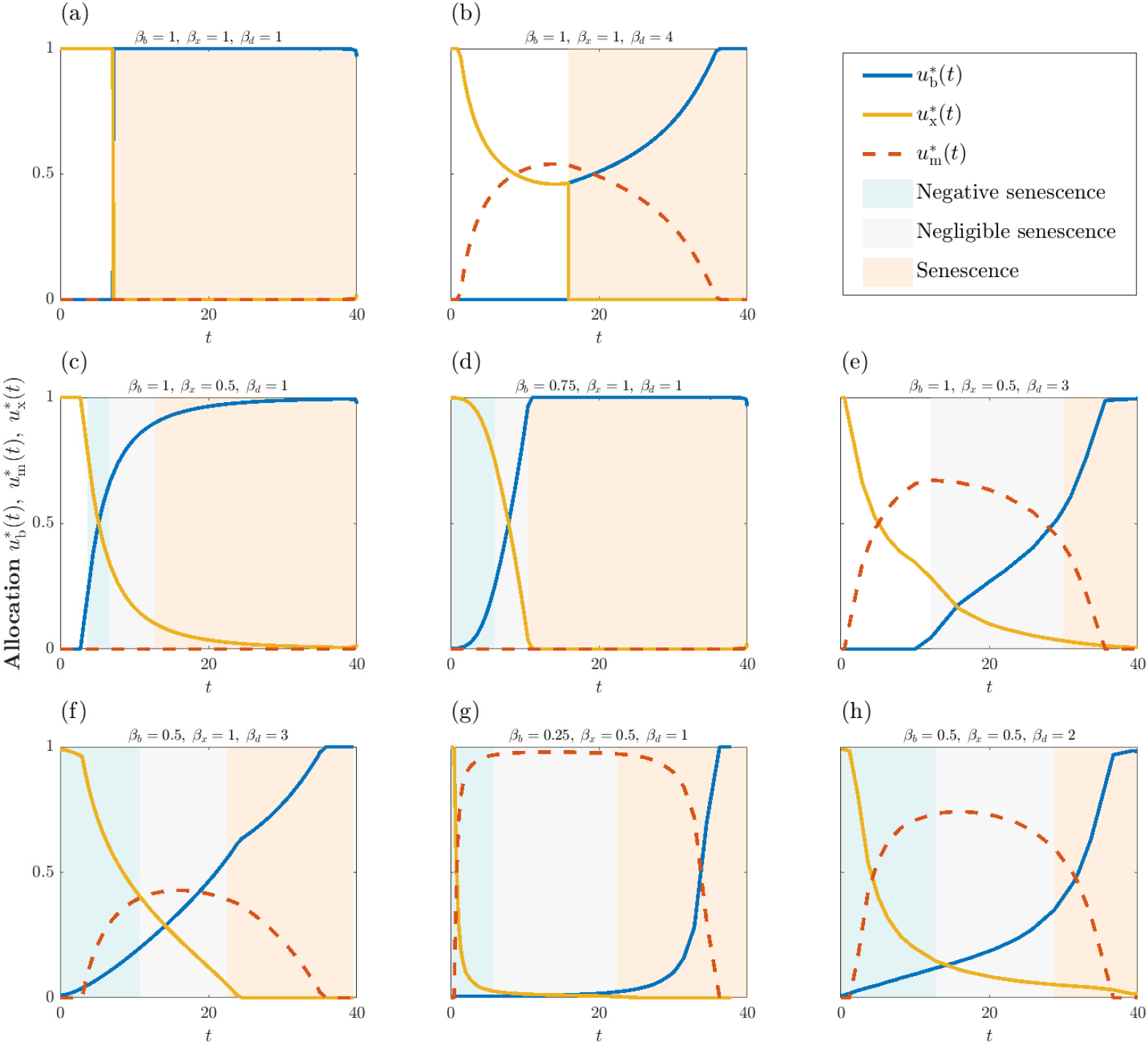
Uninvadable energy allocation schedules across eight combinations of the scaling factors *β*_*b*_, *β*_*x*_, and *β*_*d*_. Parameter values for each example listed in Table 2

In Panel (a) in Fig. 6 we set all energy allocation functions to linear scaling (*β*_*b*_ = *β*_*x*_ = *β*_*d*_ = 1), which produces a classical bang-bang allocation strategy between reproduction and growth with no maintenance on the uninvadable path. The reproductive phase is characterised by senescence.

In Panel (b) we add diminishing returns to somatic maintenance (*β*_*d*_ *>* 1), which allows interior solutions (Phases II and V) on the uninvadable path. This means that in the uninvadable path, less maintenance investment has marginally better fitness returns. The reproductive phase is again characterised by senescence. We analyse this case of determinate growth and senescence further in Section 3.1 of the main text.

In Panels (c) and (d) we illustrate cases where either growth (*β*_*x*_ *<* 1) or reproductive allocation (*β*_*b*_ *<* 1) has diminishing returns, while maintenance scales linearly (*β*_*d*_ = 1). Both produce simultaneous allocation to reproduction and growth (Phase IV), but maintenance does not appear on the uninvadable path. Allocation to reproduction increases reproductive output directly and growth allocation increases future reproductive capacity, but maintenance allocation only improves future survival. In these two cases the interior optimum is found from the trade-off between reproduction and growth, and it is never uninvadable to allocate any resources to maintenance when it scales linearly (*β*_*d*_ = 1). Growth after maturity creates short periods of negative and negligible senescence, as rapid growth momentarily reduces mortality through the size-dependent component of eq. (4). This means that there is no maintained period of non-senescence and the reproductive period is again characterised by senescence.

In Panel (e) we set two allocation functions to diminishing returns simultaneously (*β*_*x*_ *<* 1, *β*_*d*_ *>* 1). Diminishing returns to growth mean that the marginal cost of withdrawing resources from growth rises, allowing growth to continue alongside reproduction. Similarly, the marginal benefit of maintenance falls with investment, and the uninvadable path now includes simultaneous allocation to all life history functions (Phase III). Notably, the condition for negative senescence is not met and the reproductive period is characterised by negligible senescence. Transient decline in mortality after maturity is possible for other parametrisations but generally it holds that when reproduction scales linearly (*β*_*b*_ = 1) there is no “sustained” period of negative senescence. Even if growth and maintenance allocation continues after the start of reproduction.

In Panels (f) and (g) we alternately set the scaling of growth and maintenance to linear, while keeping the other two functions at diminishing returns. Both combinations produce Phase III on the uninvadable path, which means indeterminate growth with maintenance. Now a period of negative senescence can appear on the uninvadable path. Notably, maintenance does not need to have diminishing returns for it to appear on the uninvadable path, provided the other two allocation functions scale nonlinearly. In Panel (g) we set strongly diminishing returns to reproduction (*β*_*b*_ = 0.25). When maintenance scales linearly, this produces a life history in which it is uninvadable to allocate almost all of the resources to maintenance for the majority of the lifespan.

In Panel (h) we set all life history functions to diminishing returns (*β*_*b*_ = *β*_*x*_ = 0.5, *β*_*d*_ = 2). This combination produces Phase III and the reproductive period is characterised by non-senescence. We analyse this case of indeterminate growth and negative senescence further in Section 3.2 of the main text.

Across different *β*-parameter combinations, the pre-maturity phase structure, that is, whether the life history begins in Phase I (growth only) or Phase II (growth and maintenance), depends not only on the *β*-parameters but also on other model parameters such as initial size *x*_0_ and the growth return scaling *α*_*x*_. Regardless of the parameter combination, all life histories eventually converge to Phase VI (reproduction only). This is a general consequence of the life history allocation structure of the model: as survival probability declines with age, there is always a point beyond which investment in growth or maintenance no longer increases expected future fitness, and the soma becomes disposable.

### C.2 Accelerating returns to reproduction

In some cases it may be justified to depart from the more general law of diminishing returns. Accelerating returns to reproductive allocation is one such case. This requires some kind of multiplicative effect relative to the energy allocated to reproduction. Such a relationship has been proposed at least in flowering plants, where seed production has been found to increase at an accelerating rate relative to increased flowering (Schaffer and Schaffer, 1979). In that example, the synergistic effects arising from pollinator activity drive accelerating returns. We can choose to model this by setting *β*_*b*_ *>* 1. If the other two *β*-parameters are linear, this does not change the allocation strategy from the classical bang-bang outcome (Panel (a) in Fig. 6), but by adding nonlinear returns to the other two functions we again find interior solutions. Two examples are presented below. In general, the trade-off on the uninvadable path is bang-bang between reproduction and growth, but during the growth phase before reproduction there is somatic maintenance (Phase II). When reproduction begins, all allocation is directed to reproduction and the reproductive phase is characterised by rapid damage accumulation and senescence (Panel (i) in Fig. 7). However, when we increase the nonlinearities in growth and somatic maintenance, interior solutions become possible for the reproductive phase, in which small amounts of growth and maintenance are sustained for a short period after reproduction begins (Panel (j) in Fig. 7). In general, when returns to reproduction are accelerating (*β*_*b*_ *>* 1), negative senescence does not appear on the uninvadable path.

**Figure 7.**
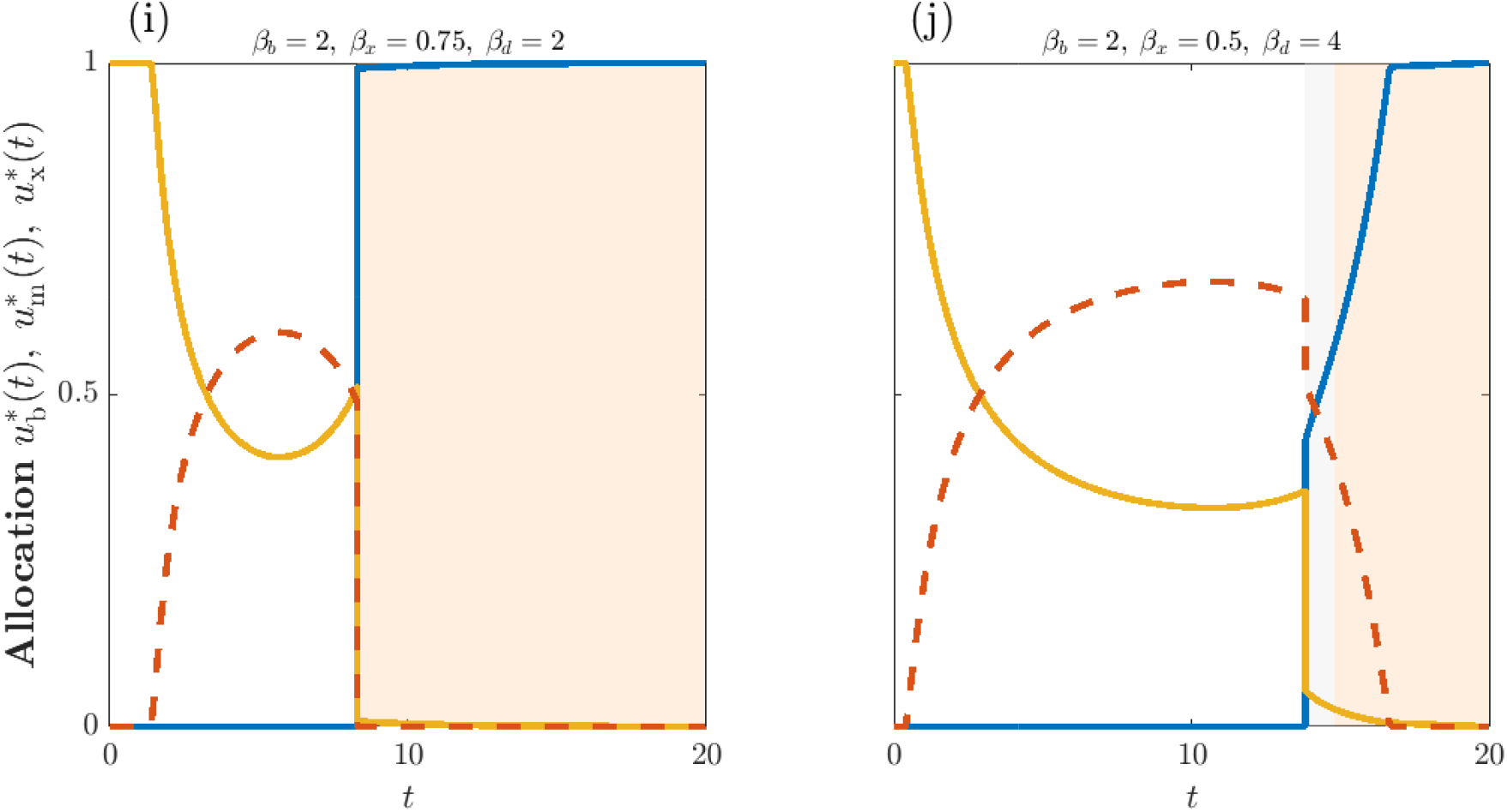
Uninvadable life history schedules for two parameter sets with accelerating returns to reproduction (*β*_*b*_ *>* 1). Parameters held constant across both panels: *a* = 1, *α*_*x*_ = 1, *c* = 0.75, *β*_*b*_ = 2, *α*_*b*_ = 0.05, *α*_*d*_ = 0.01, *µ*_e_ = 0.001, *µ*_x_ = 0.17, *κ* = −1. Panel (i): *β*_*x*_ = 0.75, *β*_*d*_ = 2. Panel (j): *β*_*x*_ = 0.5, *β*_*d*_ = 4.

